# Linking reduced prefrontal microcircuit inhibition in schizophrenia to EEG biomarkers in silico

**DOI:** 10.1101/2023.08.11.553052

**Authors:** Sana Rosanally, Frank Mazza, Heng Kang Yao, Faraz Moghbel, Hannah Seo, Etay Hay

**Author notes:** These authors contributed equally (Sana Rosanally, Frank Mazza, Heng Kang Yao).

## Abstract

Reduced cortical inhibition by parvalbumin-expressing (PV) interneurons in schizophrenia is thought to be associated with impaired cortical processing in the prefrontal cortex and altered EEG signals such as oddball mismatch negativity (MMN). Recent studies also suggest loss of somatostatin (SST) interneuron inhibition. However, establishing the link between reduced interneuron inhibition and reduced MMN experimentally in humans is currently not possible. To overcome these challenges, we simulated spiking activity and EEG during baseline and oddball response in detailed models of human prefrontal microcircuits in health and schizophrenia, with reduced PV and SST interneuron inhibition as constrained by postmortem patient data. We showed that reduced PV interneuron inhibition can account for the decreased MMN amplitude seen in schizophrenia, with a threshold below which the amplitude effect was low as seen in at-risk patients. In contrast, reduced SST interneuron inhibition did not affect the MMN amplitude. We further showed that both types of inhibition loss were necessary to account for changes in resting EEG in schizophrenia, with reduced SST interneuron inhibition increasing theta power, and reduced PV interneuron inhibition leading to a right shift from alpha to beta frequencies. Our study thus links reduced PV and SST interneuron inhibition in schizophrenia to distinct EEG biomarkers that can serve to improve stratification and early detection using non-invasive brain signals.

## Introduction

Cortical dysfunction in schizophrenia involves changes in processing across brain areas [1] and at the cellular and microcircuit level [2–5]. Altered brain activity underlying impairments in schizophrenia may also have signatures in brain signals obtained by electroencephalography (EEG) [1,6], which offer a promising source of objective and quantitative biomarkers [7] to improve patient stratification and early diagnosis, especially when symptoms are mild [8]. However, the link between cellular and microcircuit mechanisms of schizophrenia to altered cortical activity and EEG biomarkers remains to be established.

Previous studies indicate that changes in the microcircuitry of the prefrontal cortex (PFC) may underly impaired cognitive functions in schizophrenia [9–11]. Reduced inhibition has been implicated as a key mechanism, whereby post-mortem studies in schizophrenia patients found a reduced expression of parvalbumin (PV) and GAD67 in PV interneurons in PFC [2,5,12,13]. PV interneurons provide important and timely inhibition of pyramidal (Pyr) neurons [14], and modulate brain oscillations in high frequencies (gamma band, 20 – 80 Hz) [15,16] which support cognitive functions in PFC [17]. Accordingly, reduced GABA neurotransmission in the PFC has been associated with impaired function [18], and with altered brain oscillations and impaired cognition in schizophrenia [19,20]. Reduced PV expression in PV interneurons indicates a loss of functionality in these neurons, supported by the accompanying decreased GAD67 expression that directly influence synaptic inhibition, as GAD67 knock-out in PFC PV interneurons in rodents resulted in loss of inhibition from PV to Pyr neurons and consequently overexcitability of Pyr neurons [21]. Overall, reduced PV and GAD67 expression in PV interneurons in schizophrenia thus indicates a reduced synaptic inhibition from these interneurons to other neurons in the microcircuit.

Another altered mechanism of PV interneurons implicated in schizophrenia is reduced excitatory innervation via NMDA receptors. Post-mortem studies found that NMDA subunit NR2A expression was mostly absent in more than half of PV interneurons in PFC in schizophrenia [22–24], whereas there was no significant change found in other NMDA subunits NR2B-D [25]. NR2A directly affects the potency of glutamate and thus mediates excitatory activity in NMDA receptors [26]. Furthermore, rodent studies found that working memory depended on NR2A in the PFC as they mediate the majority of evoked NMDA receptor currents in layer 2/3 Pyr neurons [27].

In addition to changes in PV interneurons, recent studies also showed reduced somatostatin (SST) expression in SST interneurons in schizophrenia [5], indicating a loss of inhibition from these interneurons as well. SST interneuron mediate baseline activity [28,29] through lateral inhibition [30,31], modulate brain oscillations in low frequencies (theta and alpha band, 4 - 12 Hz) [32,33], and modulate cortical response [34]. However, reduced SST interneuron inhibition is implicated in a variety of other conditions such as aging and depression [35] and its role in schizophrenia remains unknown, whereas reduced PV interneuron inhibition has been indicated as a more specific mechanism of schizophrenia.

Altered PFC microcircuitry in schizophrenia may also underlie the respective changes seen in EEG signals [36]. In particular during auditory oddball task, which involves the presentation of a series of same-frequency tones followed by a different tone (the deviant tone or “oddball”), schizophrenia patients show a reduced performance and a smaller difference between the PFC EEG signal response to the standard and deviant tones ∼100 - 160 ms post-stimulus, referred to as mismatch negativity (MMN) [6,37]. Recordings of spike activity in monkey auditory cortex and PFC show that microcircuits in PFC process the difference between the expected and actual subsequent stimulus and generate a larger spike response to oddball stimuli, which is associated with MMN in the EEG [38]. Changes in PV or SST interneuron inhibition can lead to detectible signatures in EEG signals since these interneurons closely modulate inputs to Pyr neurons, which are the main contributors to EEG signals [39]. In addition to altered EEG during oddball response, studies found changes in resting-state EEG in schizophrenia such as increased theta and beta power vs decreased alpha power [40–44], but the underlying mechanisms remain unknown.

Whereas experimental studies either implicated cellular mechanisms of schizophrenia post-mortem or characterized changes in EEG features in living patients, the link between the two remains to be established in humans due to technical and ethical limitations in probing microcircuitry in the living human brain. Computational models offer a powerful tool for overcoming the challenges, and have been previously used to link altered ion channel mechanisms in schizophrenia to their EEG biomarkers [45]. Previous studies mainly used rodent microcircuit models to study mechanisms of EEG in health and disease [45,46], but modeling human cortical microcircuits to identify these links is motivated by several reasons. Although there are many circuit and cellular similarities between rodents and humans such as intrinsic firing properties and connectivity patterns between different cell types [47,48], there remain some important differences. Inhibitory synapses from SST and PV interneurons onto Pyr neurons are stronger in humans, have lower synaptic failures and larger postsynaptic potential (PSP) amplitudes [31,49,50]. Furthermore, there is a higher connection probability between Pyr neurons in human cortical layer 2/3 [51]. Pyr neurons are larger in humans and have longer and more complex dendrites which affect input integration [52,53]. The increased availability of human neuronal and synaptic connectivity data [31,51,54] has enabled the generation of detailed models of human cortical microcircuits [55,56], which were used to simulate microcircuit activity in health and disease as well as local microcircuit-generated EEG signals, and link changes in microcircuit mechanisms to EEG biomarkers [57].

In this study, we identified EEG biomarkers of reduced PV and SST interneuron inhibition in detailed models of human PFC microcircuits in schizophrenia, which implemented two key altered PV interneuron mechanisms and reduced SST interneuron inhibition as estimated from gene expression changes in schizophrenia. We modeled oddball response as constrained by spike recordings in monkeys in previous studies and linked reduced PV vs SST interneuron inhibition to altered PFC activity, MMN and resting EEG in schizophrenia.

## Results

We simulated baseline and oddball response activity in models of human PFC microcircuits in health and schizophrenia. We first modeled healthy PFC microcircuits by adapting our previous detailed models of human cortical L2/3 microcircuits (Fig. 1A-B) using the proportions of Pyr neurons and PV/SST/VIP interneurons in human PFC as seen experimentally (Fig. 1C). To simulate the healthy baseline activity of the PFC microcircuit, all neurons received background random excitatory input corresponding to baseline cortical and thalamic inputs. The baseline firing rates of Pyr neurons and PV interneurons in the PFC microcircuit models (Fig. 1D) were within the range measured *in vivo* in humans, Pyr: 0.73 ± 0.03 Hz (experimental: 0.66 ± 0.51 Hz), PV: 6.64 ± 0.17 Hz (experimental: 2.63 ± 2.55 Hz). Baseline firing rates of SST and VIP interneurons were on the order of magnitude seen in rodents, SST: 3.64 ± 0.14 Hz (experimental: 6.3 ± 0.6 Hz), VIP: 2.51 ± 0.14 Hz (experimental: 3.7 ± 0.7 Hz). We therefore left the rest of the microcircuit parameters unaltered from the previous models. The power spectral density (PSD) of the simulated EEG from the microcircuit models during baseline activity exhibited a peak in the theta (4 – 8 Hz) and alpha (8 – 12 Hz) frequency bands (Fig. 1E), as well as a 1/f relationship, all of which were in line with spectral properties of human prefrontal resting-state EEG activity.

**Figure 1.**
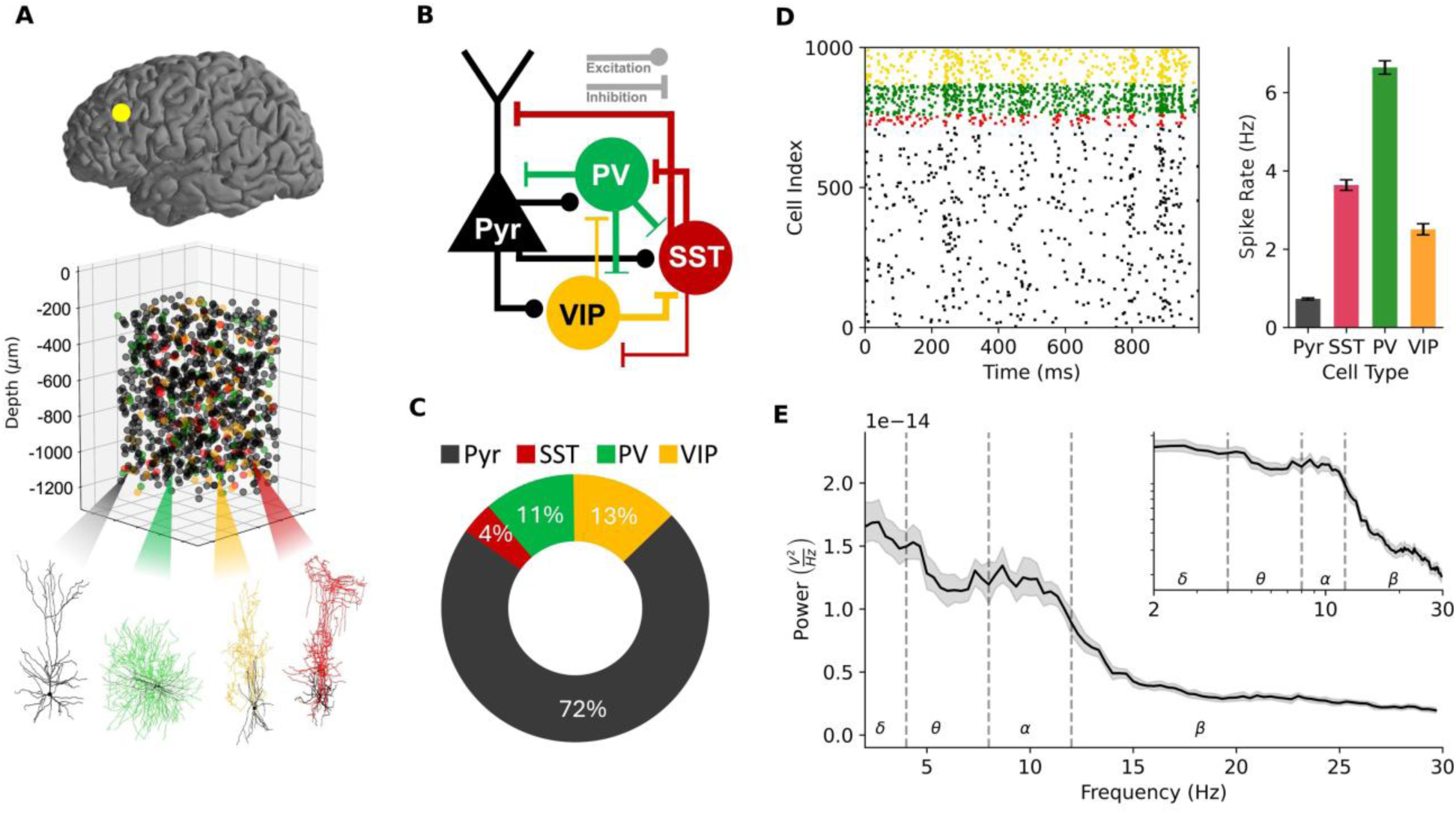
Detailed models of human prefrontal microcircuit baseline spiking activity and EEG. **A.** Detailed models of human PFC microcircuits showing placement of 1000 connected neurons, human neuronal morphologies of the four key neuron types (green: PV, red: SST, black; Pyr, yellow: VIP). **B.** Connectivity diagram between neuron types in the microcircuit. **C.** Cellular proportions of each neuron type: Pyr (72%), SST (4%), PV (11%), VIP (13%). **D.** Left - raster plot of neuronal spiking in the microcircuit at baseline, color-coded according to each neuron type. Right - baseline firing rates of all neurons (mean and SD, n = 30 random microcircuits). **E.** PSD plot of simulated EEG at baseline (mean across n = 30 random microcircuits, showing bootstrap mean and 95% confidence interval. Canonical frequency bands are shown by vertical dotted lines. Inset: the same plot in log-log scale, showing 1/f relationship.

We modelled microcircuit changes in schizophrenia by implementing two key mechanisms involving PV interneuron inhibition (Fig. 2A) according to human PFC post-mortem studies. The first mechanism (referred to as the output mechanism) was a reduced PV interneuron synaptic and tonic inhibition conductance, and the second mechanism (referred to as the input mechanism) was a reduction in NMDA synaptic conductance from Pyr neurons to PV interneurons. We simulated baseline activity, which in the PFC also corresponded to the activity during presentation of standard tones, and simulated oddball (deviant tone) response in the healthy and schizophrenia microcircuits.

**Figure 2.**
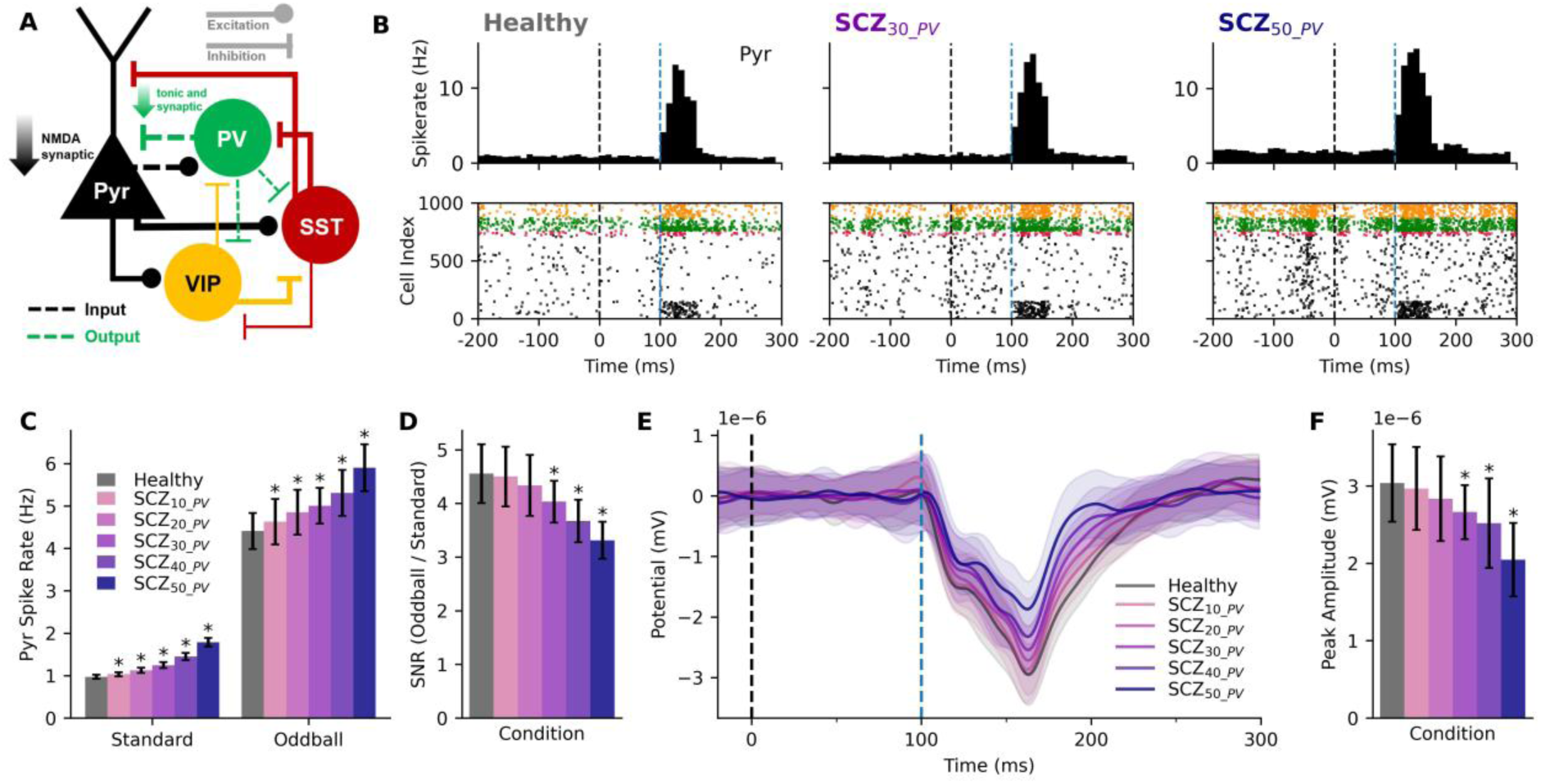
Simulated oddball response and MMN in healthy and schizophrenia microcircuits. **A.** Connectivity diagram of the schizophrenia microcircuit models showing the altered PV output and input mechanisms (green and black dashed lines). **B.** Peristimulus time histogram (top, n = 50 randomized microcircuits) and raster plot (bottom) of simulated oddball response in a healthy microcircuit (left) and schizophrenia microcircuits with 30% (middle) or 50% (right) reduced PV interneuron inhibition. Healthy models reproduced response firing rates and profiles from previous experimental studies in primates. Black dashed line denotes the external stimulus time and blue dashed line denotes PFC activation time. **C.** Average Pyr spike rate during simulated standard (baseline) and oddball response in healthy and schizophrenia microcircuits with 10 - 50% reduced inhibition (mean and SD). **D.** SNR in healthy and schizophrenia microcircuits. **E.** Difference between simulated ERP during oddball vs standard response in healthy and schizophrenia microcircuit models (oddball - standard response, mean and SD, n = 50 randomized microcircuits). **F.** Peak amplitude of MMN response in healthy and schizophrenia microcircuits. Asterisks in parts C, D, F indicate significance p < 0.05.

We modeled healthy oddball response by stimulating a population of Pyr neurons and PV and SST interneurons to reproduce the firing rate profile of Pyr neurons along the period 100 – 160 ms post-stimulus as recorded in primates (Fig. 2B). We then applied the same stimulus paradigm to schizophrenia microcircuit models with 10 - 50% reduced PV interneuron inhibition, while also applying the respective reduced NMDA mechanism effect on the stimulus synapses onto PV interneurons.

Both baseline and oddball response firing rates increased linearly and moderately with reduced PV interneuron inhibition, with a supralinear jump in effect for 50% reduction (Fig 2C). The effect was larger for baseline activity (healthy: 0.97 ± 0.05 Hz; SCZ_30_: 1.24 ± 0.07 Hz, +28%, *p* < 0.05, *d* = 4.46; SCZ_50_: 1.79 ± 0.10 Hz, +84%, *p* < 0.05, *d* = 10.05) compared to oddball response (healthy: 4.41 ± 0.43 Hz; SCZ_30_: 5.0 ± 0.42 Hz, +13% *p* < 0.05, *d* = 0.9; SCZ_50_: 5.9 ± 0.56 Hz, +34%, *p* < 0.05, *d* = 2.98). Consequently, the signal-to-noise ratio (SNR) of firing rates during oddball response (signal) vs standard response (noise) decreased with inhibition reduction (Fig 2D, healthy: 4.56 ± 0.55; SCZ_30_: 4.03 ± 0.39, -11%, *p* < 0.05, *d* = -1.09; SCZ_50_: 3.31 ± 0.35, -27%, *p* < 0.05, *d* = -2.68).

To determine the effect of reduced PV interneuron inhibition on the MMN amplitude of the ERP during oddball response, we simulated the EEG from the microcircuits during baseline and oddball response and plotted the difference (Fig. 2E). The MMN amplitude in schizophrenia microcircuits decreased linearly and moderately with inhibition reduction, with a supralinear jump in effect for 50% reduction (Fig. 2E,F; healthy: 3.04 ± 0.50 nV; SCZ_30_: 2.66 ± 0.35 nV, -12%, *p* < 0.05, *d* = -0.87; SCZ_40_: 2.52 ± 0.58 nV, -17%, *p* < 0.05, *d* = -0.95; SCZ_50_: 2.05 ± 0.48 nV, - 33%, *p* < 0.05, *d* = -2.01).

We next compared the effects of reduced PV and SST interneuron inhibition, by simulating schizophrenia microcircuits with either 40% reduced PV or SST interneuron inhibition (SCZ_40_PV_ or SCZ_40_SST_, respectively) or both PV and SST interneurons (SCZ_40_PV+SST_). Baseline Pyr neuron firing rates (Fig. 3A) increased to a similar extent in microcircuits with reduced PV or SST interneuron inhibition (SCZ_40_PV_: +48%, 1.39 ± 0.08 Hz, *p* < 0.0005, Cohen’s *d* = 7.2; SCZ_40_SST_: +32%, 1.25 ± 0.06 Hz, *p* < 0.0005, Cohen’s *d* = 5.7). The effect in SCZ_40_PV+SST_ was larger than the sum of the separate effects, and was rather multiplicative (+104%, 1.93 ± 0.10 Hz, *p* < 0.0005, Cohen’s *d* = 12.1). Oddball response increased in SCZ_40_PV_ (+19%, 5.36 ± 0.54 Hz, *p* < 0.0005, Cohen’s *d* = 1.8), with a minor effect in SCZ_40_SST_ (-6%, 4.22 ± 0.43 Hz, *p* = 0.002, Cohen’s *d* = 0.6). The combined effect in SCZ_40_PV+SST_ was mostly equivalent to the effect in reduced PV interneuron inhibition alone (+24%, 5.55 ± 0.52 Hz, *p* < 0.0005, Cohen’s *d* = 2.2). The SNR decreased in all schizophrenia microcircuits, with a larger effect for SCZ_40_SST_ (-29%, 3.39 ± 0.37, *p* < 0.0005, Cohen’s *d* = 2.9, Fig. 3A) compared to SCZ_40_PV_ (-19%, 3.85 ± 0.41, *p* < 0.0005, Cohen’s *d* = 1.9), and the combined effect in SCZ_40_PV+SST_ was less than the sum of the separate effects (-40%, 2.89 ± 0.30, *p* < 0.0005, Cohen’s *d* = 4.2).

**Figure 3.**
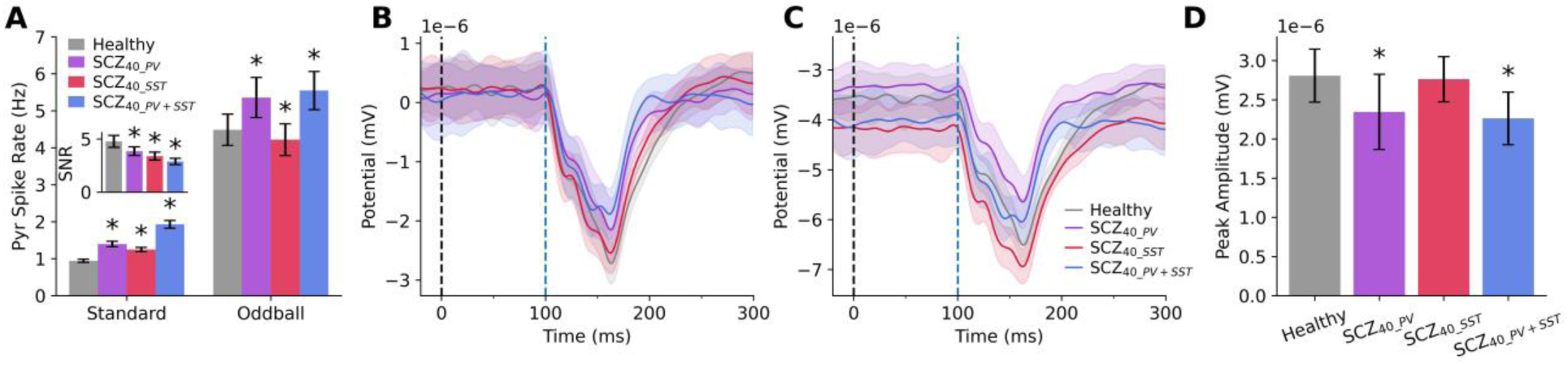
Effects of PV vs SST interneuron inhibition on oddball response and MMN. **A.** Pyr spike rate during simulated standard and oddball response in healthy and schizophrenia microcircuits with either 40% reduced PV (purple) or SST (red) interneuron inhibition or both (blue; n = 50 randomized microcircuits per condition). Inset: SNR of Pyr neurons in the different conditions. **B.** Difference between simulated ERP during oddball vs standard response in healthy and the different schizophrenia microcircuit models (oddball - standard response). Black and blue dashed lines denote stimulus and PFC activation times, respectively. **C.** Simulated ERP during response to oddball alone. **D.** Peak amplitude of MMN response in healthy and schizophrenia. All plots show mean and SD. Asterisks indicate significance *p* < 0.05.

Interestingly, schizophrenia microcircuits with reduced SST interneuron inhibition did not have an altered MMN compared to healthy (SCZ_40_SST_: 2.76 ± 0.29 pV, *p* = 0.471, Cohen’s *d* = 0.1, Fig. 3B, D). Even when combined with reduced PV interneuron inhibition, the effect did not differ (*p* = 0.32) from that seen in microcircuit with reduced PV interneuron inhibition alone (SCZ_40_PV+SST_: -19%, 2.26 ± 0.34 pV, *p* < 0.0005, Cohen’s *d* = 1.6; SCZ_40_PV_: -16%, 2.35 ± 0.48 pV, *p* < 0.0005, Cohen’s *d* = 1.1). The reason for this was because SST affected both baseline (-14%, *p* < 0.0005, Cohen’s *d* = 0.9, Fig. 3C) and oddball ERP (-9%, *p* < 0.0005, Cohen’s *d* = 0.8), and the changes were cancelled out in the difference waveform used to measure MMN amplitude.

We next characterized signatures of the altered inhibition effects on simulated resting-state EEG by comparing the PSD in healthy and schizophrenia microcircuit models (Fig. 4). Simulated EEG from schizophrenia microcircuits with reduced PV interneuron inhibition showed a prominent peak in the alpha band (8 - 12 Hz) as the healthy microcircuit models but exhibited a rightward shift (Fig. 4A). We decomposed the EEG PSD into aperiodic (Fig. 4B) and periodic (Fig. 4C) components to compare the distinct functional components of PSD. There were no major changes in aperiodic broadband power (Fig. 4B), but there was a large rightward shift in periodic peak alpha frequency from 10.1 to 12.4 Hz in SCZ_40_PV_ compared to healthy (+23%, *p* < 0.0005, Cohen’s *d* = 1.5, Fig. 4C), evident also in a large increase in low beta (12 – 20 Hz) power compared to healthy (+63%, *p* < 0.0005, Cohen’s *d* = 4.78). There was also a small rightward shift in peak alpha frequency from 10.1 to 11.2 Hz in SCZ_20_PV_ compared to healthy (+11%, *p* = 0.001, Cohen’s *d* = 0.8).

**Figure 4.**
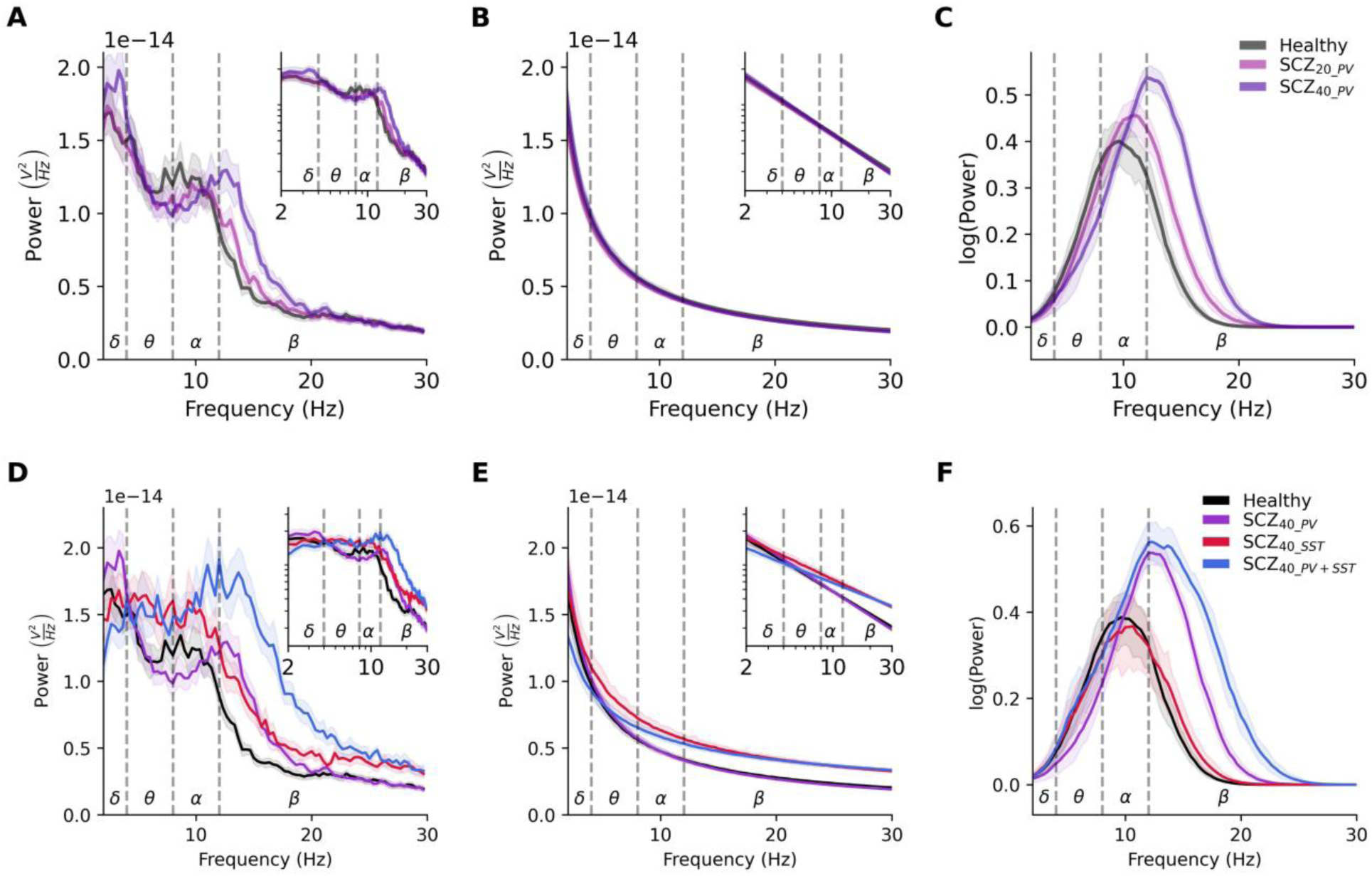
Resting-state EEG signatures of reduced PV vs SST interneuron inhibition in schizophrenia microcircuit models. **A.** PSD of simulated EEG from the healthy (black) and schizophrenia microcircuit models with 20% (light purple) and 40% (dark purple) reduced PV interneuron inhibition (n = 30 randomized microcircuits per condition, bootstrapped mean and 95% confidence interval) Insets show the same PSD on the log-log scale. Dashed lines delimit the frequency bands. **B-C.** Aperiodic (B) and periodic (C) components of the PSD in healthy and schizophrenia microcircuits with reduced PV interneuron inhibition. **D.** PSD of simulated resting-state EEG from healthy (black) and schizophrenia microcircuit models with either 40% reduced PV (purple) or SST (red) interneuron inhibition or both (blue; n = 25 randomized microcircuits per condition, bootstrapped mean). **E-F.** Aperiodic (E) and periodic (F) components of the PSD in the healthy and schizophrenia microcircuits with reduced PV vs SST interneuron inhibition.

In contrast, in microcircuits with reduced SST interneuron inhibition there was increased power in all bands (Fig. 4D, SCZ_40_SST_: theta: +23%, *p* < 0.0005, Cohen’s *d* = 2.1; alpha: +25%, *p* < 0.0005, Cohen’s *d* = 2.0; beta: +60%, *p* < 0.0005, Cohen’s *d* = 5.2). SCZ_40_PV+SST_ microcircuits had altered PSD that combined the effects of PV and SST interneuron inhibition reduction, with increased theta power (+17%, *p* < 0.0005, Cohen’s *d* = 1.7), alpha power (+36%, *p* < 0.0005, Cohen’s *d* = 2.9) and beta power (+187%, *p* < 0.0005, Cohen’s *d* = 10.9), and a right shift in peak frequency from alpha to beta, though also a decrease in absolute power in delta (-23%, *p* < 0.0005, Cohen’s *d* = 2.0). Unlike SCZ_40_PV_ microcircuits which showed no change in aperiodic features, SCZ_40_SST_ microcircuits showed a decrease in aperiodic exponent (Fig. 4E, -22%, *p* < 0.0005, Cohen’s *d* = 2.0) and an increase in broadband power (+29%, *p* < 0.0005, Cohen’s *d* = 1.5). SCZ_40_PV+SST_ microcircuits exhibited a similar decrease in exponent (-35%, *p* < 0.0005, Cohen’s *d* = 3.4) and increase in broadband power (+17%, *p* < 0.0005, Cohen’s *d* = 1.3), but also showed a decrease in offset (-34%, *p* < 0.0005, Cohen’s *d* = 1.6). Reduced SST interneuron inhibition had no effect on the periodic component (Fig. 4E), and SCZ_40_PV+SST_ microcircuits had a similar right shift in peak alpha frequency as seen in SCZ_40_PV_ microcircuits.

## Discussion

Our study mechanistically links altered cell-specific inhibition with clinically relevant EEG changes in schizophrenia. Using detailed simulations of baseline activity and oddball response of human prefrontal microcircuits constrained by human cellular, circuit and gene-expression data in health and schizophrenia, we showed that reduced PV interneuron inhibition in schizophrenia can account for the decrease in MMN amplitude seen in patients, whereas reduced SST interneuron inhibition effects were limited to resting EEG. Our results thus establish PV and SST interneuron inhibition as target mechanisms for new treatments and indicate a primary role of PV interneuron inhibition as an underlying mechanism in schizophrenia. Moreover, our results show a threshold effect of reduced inhibition and make testable quantitative predictions about the degree of reduced inhibition that underlies decreased MMN amplitude in different severities, which may improve the subtyping of schizophrenia and early detection of the disorder when it is still largely asymptomatic.

Our results establish a primary role of reduced PV interneuron inhibition as an underlying mechanism of the decreased MMN amplitude seen in patients [58,59], and thus validate previous hypotheses [60]. The lack of effect of reduced SST interneuron inhibition on MMN is supported by the role of these interneurons in maintaining baseline activity [28,29], which in our simulations led to changes in both baseline ERP and oddball ERP that cancelled each other. Interestingly, the reduction in MMN amplitude had a threshold effect, with a 50% inhibition reduction leading to 33% decrease in MMN, within the range seen in schizophrenia patients [58,59]. Smaller levels of reduced inhibition (10 - 40%) led to a milder effect on MMN which correspond well with the 15% decrease seen on average in individuals at clinically high risk for schizophrenia [8,61]. Our results thus suggest that the at-risk population may already have a reduced PV interneuron inhibition, which may underlie the milder symptoms that later develop into more severe symptoms in schizophrenia [61]. Thus, our models of schizophrenia PFC microcircuits characterize the implications of different levels of reduced PV interneuron inhibition on the MMN response, which enable a mechanistic stratification and improved early detection and outcome prediction in at-risk population [59].

Although reduced SST interneuron inhibition on its own did not affect the MMN amplitude, the decreased MMN amplitude in schizophrenia microcircuits was mediated by the reduced PV interneuron inhibition disinhibiting SST interneurons, consequently enhancing their inhibition of the distal apical dendrites of Pyr neurons [30]. The apical dendrites generally act as a negative current source by receiving a large number of excitatory inputs, and as the distance between the apical dendrites source and the soma and basal dendrites sink is large, their difference in potential is the major contributor of the dipole from the neuron. Increased SST interneuron inhibition of the apical dendrites due to the reduced PV interneuron inhibition makes the apical dendrites less positively charged, which consequently reduces the electric dipole of the neuron and thus the MMN amplitude measured by EEG [62].

Reduced inhibition also accounted well for changes seen in resting state EEG in schizophrenia. The rightward shift in the peak frequency from high-alpha band to low-beta band due to reduced PV interneuron inhibition in our simulated schizophrenia microcircuits is a key change seen in schizophrenia patients [40–44]. Reduced power in alpha frequency band has been associated with schizophrenia positive and negative symptoms and with chronicity [63], whereas increased beta power is associated with increased distraction [64,65] and is a symptom of schizophrenia. Reduced SST interneuron inhibition was necessary to reproduce the increased theta power seen in schizophrenia patients [44], in agreement with previous studies that showed the role of SST interneurons in modulating low frequencies [33].

Our MMN and resting-state EEG biomarkers can be applied on patient data, with simulation of additional levels of PV and SST interneuron inhibition reduction, to better stratify schizophrenia patients and facilitate early detection in at-risk population. To that effect, machine learning methods can be trained to estimate the mechanisms from in-silico data using normalized biomarkers [66] that can be applied to patient data, or the simulated EEG can be scaled to account for the difference between the number of neurons in our models compared to the amount of neurons that generate the recorded EEG at a given electrode, which would be about a factor of 10,000 to 100,000 [67,68].

The reduced SNR of cortical processing in our schizophrenia microcircuit models was mainly driven by increased baseline firing rates resulting from the reduced PV and SST interneuron inhibition. The increase in baseline rates is in agreement with previous fMRI and EEG studies that showed increased baseline activity in schizophrenia, associated with positive symptoms [69,70].

The link between gene expression and inhibitory interneuron synaptic function that we implemented in our models is not trivial, but a few factors support our framework. The reduced PV expression was accompanied by reduced GAD67, an enzyme that synthesizes GABA and directly affects synaptic inhibition [71]. In living rodents, a reduction of cortical PV interneurons elicited deficits in social behaviour and cognition relevant to schizophrenia symptoms [72]. The change in gene expression could alternatively correspond to a reduced number of synapses [73], however the net decrease in inhibition would be mostly similar. We also simplified the implementation of the 50% reduction in NMDA NR2A subunit, which is expressed in 40% of the PV interneurons in PFC [24], as an overall 20% reduction in synaptic NMDA conductance across all connections between Pyr and PV interneurons in the microcircuit, assuming that the net decrease in inhibition would be mostly similar in either case. We found that the reduction in MMN amplitude seen experimentally in schizophrenia corresponded to a larger inhibition reduction as suggested by the percent change in expression, suggesting that the percent change in gene expression may translate to a larger physiological effect in terms of NMDA and synaptic conductance. Relatedly, bulk-tissue expression studies indicate that some layers (3 and 4) may involve a larger reduction of PV expression than the average expression across L2/3 [2] that we implemented and thus could underlie the larger physiological effect.

Our models were constrained with microcircuit data taken from PFC where possible, and various other brain regions due to limited availability of human neuronal and microcircuit data, thus the models aimed to primarily represent canonical cortical microcircuits. While different brain regions such as the PFC and sensory regions have some differences in wiring and function when taking into account all six layers of the cortex, layers 2 and 3 microcircuitry (which was the focus of our models) is similar across regions [74]. Finally, for this study we modelled oddball processing in a single region (PFC microcircuit) rather than the interaction between multiple brain regions. While oddball processing may involve ongoing interactions between PFC and other brain regions such as the primary and secondary auditory cortex [75], the main computation of the MMN ignal is performed in the PFC [38,75], thus supporting modeling and studying it in isolation. However, future studies can simulate multiple microcircuits to study the multi-regional aspects of oddball processing in health and schizophrenia.

## Methods

### PFC microcircuit models

We modeled detailed human PFC microcircuit models by adapting our previous models of human cortical layer 2/3 microcircuits [55], which consisted of 1000 connected neurons distributed randomly in a 500x500x950 μm^3^ volume, situated 250µm - 1200μm below the pia (corresponding to human L2/3) [53]. The model included four key neuron types: Pyr neurons, SST interneurons, PV interneurons, and VIP interneurons, with detailed reconstructed human morphologies. To adapt the models to the human PFC, we applied the neuron type proportions as measured in human PFC tissue, with 72% Pyr neurons and 28% interneurons [76]: 11% PV, 4% SST and 13% VIP interneurons. Other parameters were unchanged from the previous models (see supplementary tables in Yao et al 2022 [55] for synaptic conductance and probability of the different connections, and ion channel density values in the neuron models). The neuron models included sodium, potassium and calcium channels in the axon and soma, and h-current in the dendrites, and reproduced the firing, dendritic sag current, and synaptic properties (amplitude and short-term dynamics) measured in human neurons. Connection probability between Pyr neurons was 0.15, as measured in humans [51], and other connection probabilities were constrained using rodent data. PV interneurons targeted the basal dendrites of Pyr neurons, whereas SST interneurons targeted the apical dendrites of Pyr neurons. Excitatory synapses included AMPA and NMDA (*τ_rise,NMDA_* = 2 ms; *τ_decay,NMDA_* = 65 ms; *τ_rise,AMPA_* = 0.3 ms; *τ_decay,AMPA_* = 3 ms), whereas inhibitory synapses were GABA_A_ (*τ_rise,GABA_* = 1 ms; *τ_decay,GABA_* = 10 ms). The synaptic reversal potential were *E_exc_* = 0 mV and *E_inh_* = -80 mV. We modelled tonic inhibition using a model for outward rectifying tonic inhibition [77], with G_tonic_ = 0.938 mS/cm^2^ for all neurons. The models were simulated using NEURON [78], LFPy [79] and parallel computing in high-performance grids (SciNet) [80,81].

### PFC baseline activity simulations

The microcircuit was injected with background excitatory input, simulated using Ornstein-Uhlenbeck (OU) processes [82] at each dendritic midpoint, to ensure similar levels of inputs along the dendritic path to the soma. We placed 5 additional OU processes along the apical trunk of the Pyr neuron models at 10%, 30%, 50%, 70% and 90% of the apical dendrite length. The base excitatory OU conductance was 28 pS, 280 pS, 30 pS and 66 pS for Pyr, PV, SST and VIP neurons respectively. We did not use inhibitory OU because the model microcircuit provided sufficient inhibition. We scaled the OU conductance values to increase with distance from the soma by multiplying them with the exponent of the relative distance from the soma (ranging from 0 to 1): ḡ_OU_ = ḡ ⋅ exp(X_relative_). We compared the baseline firing rates to *in-vivo* data recorded in humans (Pyr: 0.66 ± 0.51 Hz, PV: 2.63 ± 2.55 Hz) [83] or rodents (SST: 6.3 ± 0.6 Hz, VIP: 3.7 ± 0.7 Hz) [29].

### Schizophrenia microcircuit models

Schizophrenia microcircuits were modelled by altering two types of mechanisms affecting the output and input of PV interneurons. The output mechanism corresponded to a 22% reduction in PV expression in schizophrenia [2], which was modelled by a 22% reduction in synaptic and tonic inhibition from PV interneurons onto other neurons in the microcircuit. The input mechanism corresponded to a 20% decrease in NMDA subunit expression (NR2A) [24], and was modelled by a 20% reduction in NMDA synaptic conductance from Pyr neurons onto PV interneurons. We referred to these schizophrenia microcircuit models as SCZ_20_, and in addition simulated additional levels of the effect (10, 30, 40, 50%). We additionally modeled reduced SST interneuron synaptic and tonic inhibition, in line with a similar reduced expression in schizophrenia measured postmortem [5].

### Oddball response models

We simulated oddball response using the response firing profile of single neurons in monkey PFC during an oddball auditory task [38]. The experimental response spanned 100 – 160 ms post-stimulus, and the firing response profile had three phases: an increase in firing rate to 6 Hz over 100 - 120 ms, a further increase to 11 Hz over 120 – 140 ms, and a decrease to 7 Hz over 140 – 160 ms. To reproduce the experimental response, we stimulated a population 150 Pyr neurons, according to the experimental estimate of 20% responsive neurons [38]. Adapting our previous method [55], we used three activation phases of varying stimulus strength of 5 synapses, applied at a uniformly random time within the activation period to each Pyr neuron (1.3 nS during t = 97 – 117 ms, 2.75 nS during 117 – 137 ms, and 1.4 nS during 137 - 152 ms). We also stimulated 40 PV interneurons, each with 5 synapses of 2.5 nS every 10 ms between t = 95 – 155 ms, thus starting 2 ms before the Pyr neuron activation. In addition, we stimulated 30 SST interneurons, each with 5 synapses of 2.5 nS every 10 ms between t = 110 – 130 ms. This balanced excitation/inhibition activation was necessary to achieve the target firing rates, whereas activating Pyr neurons alone resulted in either too low or too high firing rates. For the PFC response to standard tones we used the baseline activity, since the experimental response to standard tones was not significantly different from baseline [38].

### Simulated microcircuit EEG

We simulated dipole moments together with neuronal spiking activity using LFPy. EEG time series was generated by the microcircuit models using the same methodologies as in previous work [57]. Specifically, we used a four-sphere volume conductor model corresponding to the brain (grey and white matter), cerebrospinal fluid, skull, and scalp with radii of 79 mm, 80 mm, 85 mm, and 90 mm, respectively, that assumes a homogeneous, isotropic, and linear (frequency-independent) conductivity. The conductivity for each sphere was 0.47 S/m, 1.71 S/m, 0.02 S/m, and 0.41 S/m, respectively [84]. The EEG power spectral density was calculated using Welch’s method [85] for 3 – 30 Hz using the SciPy python package.

### Simulated EEG event related potentials (ERP)

To analyze ERP during the oddball MMN response, simulated EEG timeseries were first lowpass filtered to 40 Hz and downsampled to 100 Hz. They were then baseline corrected using the period 0 ms - 500 ms before the stimulus. ERP potential was identified as the largest negative peak from 100 ms – 200 ms post stimulus.

### EEG periodic and aperiodic components

We decomposed EEG PSDs into periodic and aperiodic (broadband) components using tools from the FOOOF library [86]. The aperiodic component of the PSD was a 1/f function, defined by a vertical offset and exponent parameter. The periodic components were derived using up to four Gaussians, defined by center frequency (mean), bandwidth (variance) and power (height).

### Statistical tests

Statistical significance was determined wherever appropriate using two-sample t-tests. Cohen’s *d* was calculated to determine effect size.

### Data availability

All models and original code for simulations and figures have been deposited on Zenodo and will be publicly available as of the date of publication.

## Acknowledgements

This study was supported by the Krembil Foundation, by Ontario Graduate Scholarship, by a stipend award from the Department of Physiology at University of Toronto, and by Miner’s Lamp Innovation Fund.

## Competing interests

The authors declare no competing financial or non-financial interests.

## Author contributions

SR and EH contributed to conception of the work. SR, FrM, HKY and EH contributed to the design of the work. SR, FrM, HKY, HS and FaM contributed to the acquisition of data. SR, FrM, HKY, HS, FaM and EH contributed to the analysis of data. SR, FrM, HKY, HS, FaM and EH contributed to the interpretation of data. SR and EH contributed to drafting the manuscript. SR, FrM, HS, FaM and EH contributed to revising the manuscript.

## Notes

### Competing Interest Statement

The authors have declared no competing interest.

### Summary of Updates

Revised manuscript throughout after peer-review and added contributing authors

